# Metabolomic Analysis of Murine Tissues Infected with *Brucella melitensis*

**DOI:** 10.1101/2024.11.16.623915

**Authors:** Bárbara Ponzilacqua-Silva, Alexis S. Dadelahi, Charles R. Moley, Mostafa F.N. Abushahba, Jerod A. Skyberg

## Abstract

*Brucella* is a gram negative, facultative, intracellular bacterial pathogen that constitutes a substantial threat to human and animal health. *Brucella* can replicate in a variety of tissues and can induce immune responses that alter host metabolite availability. Here, mice were infected with *B. melitensis* and murine spleens, livers, and female reproductive tracts were analyzed by GC-MS to determine tissue-specific metabolic changes at one-, two- and four-weeks post infection. The most remarkable changes were observed at two-weeks post-infection when relative to uninfected tissues, 42 of 329 detected metabolites in reproductive tracts were significantly altered by *Brucella* infection, while in spleens and livers, 68/205 and 139/330 metabolites were significantly changed, respectively. Several of the altered metabolites in host tissues were linked to the GABA shunt and glutaminolysis. Treatment of macrophages with GABA did not alter control of *B. melitensis* infection, and deletion of the putative GABA transporter BMEI0265 did not alter *B. melitensis* virulence. While glutaminolysis inhibition did not affect control of *B. melitensis* in macrophages, glutaminolysis was required for macrophage IL-1β production in response to *B. melitensis*. In sum, these results indicate that *Brucella* infection alters host tissue metabolism and that these changes could have effects on inflammation and the outcome of infection.

## INTRODUCTION

Brucellosis is a worldwide bacterial zoonosis with a broad host range of animals, spanning from wildlife to agriculturally important livestock (1). The three main species recognized as substantial health threats and most common human pathogens are *B. melitensis, B. abortus*, and *B. suis* (2). Humans acquire brucellosis through inhalation of infectious aerosols and by consumption of contaminated animal products (3–5). Human brucellosis can be a severely debilitating disease that requires hospitalization (6). Human disease is characterized by persistent waves of fever with systemic symptoms that can vary among individuals, including chills, malaise, headaches, and hepato- or splenomegaly (7).

*Brucella* replicates intracellularly and has a predilection for organs rich in reticuloendothelial cells such as the spleen and liver (8). Controlling *Brucella* infection is complicated due to the fact that humoral immunity and some antibiotics are unable to reach the bacteria while inside the cell (9). Thus, antibiotic therapy is lengthened, increasing the likelihood of treatment-related side effects and leading to non-compliance among those being treated (10,11). Aggravating the situation, no vaccine is licensed to prevent human brucellosis.

Metabolic changes in immune cells have an influence on the effectiveness of the immune response and how intracellular bacteria experience stress (12–15). In the context of brucellosis, macrophages are important for controlling infection but can also be the major niche for *Brucella* survival and replication depending on their metabolic status (1). Metabolic reprogramming is associated with pathways such as glycolysis, TCA cycle, glutaminolysis, gamma-aminobutyric acid (GABA) shunt, and others, depending on the immune response (16). For example, M1 polarization of macrophages is marked by increased glycolysis and an impaired TCA cycle, driving the accumulation of several metabolites as well as promoting immune signaling and, in some cases, antimicrobial activities (17–19). In contrast, M2 polarization of macrophages is marked by mitochondrial oxidative metabolism of fatty acids, leading to an anti-inflammatory and pro-resolution profile which can favor *Brucella* replication (20,21).

It is largely unknown what metabolic conditions *Brucella* encounter across host tissues over the course of infection. In this study, we performed global screening of tissue metabolites in multiple tissues at various timepoints after *Brucella* infection. In addition, we investigated the effects of these metabolic changes on inflammatory cytokine production and the requirements for *Brucella* virulence.

## Materials and Methods

### Bacterial strains and growth conditions

Experiments with live *Brucella melitensis* were performed in biosafety level 3 (BSL3) facilities. *B. mellitensis* 16M was obtained from Montana State University (Bozeman, Montana) and grown on Brucella agar (Becton, Dickinson) for 3 days at 37°C/5% CO_2_. Colonies were then transferred to Brucella broth (Bb; Becton, Dickinson) and grown at 37°C with shaking overnight. The overnight *Brucella* concentration was estimated by measuring the optical density (OD) at 600 nm, and the inoculum was prepared and diluted to the appropriate concentration in sterile phosphate-buffered saline (sPBS). Titer was confirmed by serial dilution of the *B. melitensis* inoculum onto Brucella agar plates.

### Generation of B. melitensisΔbmeI0265

The BMEI0265 gene in *B. melitensis* 16M was replaced in frame with a chloramphenicol resistance gene (*catR*) from plasmid pKD3 (22) using the suicide plasmid pNTPS139 (23). Approximately 1,000-bp fragments upstream and downstream of BMEI0265 were amplified by PCR using primers shown in Table S1. We also generated a 1,044-bp fragment containing the *catR* gene from pKD3 using primers listed in Table S1. The 5’ end of the forward primer used to amplify the upstream fragment of BMEI0265 contained homology to 30 bp upstream of the BamHI site in pNTPS139. The 5’ end of the forward primer used to amplify the *catR* gene from pKD3 contained 30 bp homologous to the 3’ end of the upstream BMEI0265 fragment. The downstream fragment of BMEI0265 was amplified using a forward primer whose 5’ end contained 30 bp homologous to the 3’ end of the *catR* fragment, while the 5’ end of the downstream BMEI0265 fragment reverse primer contained 30 bp homologous to the 30bp downstream of the SalI site of pNTPS139. pNTPS139 was digested with BamHI/SalI. The upstream and downstream BMEI0265 fragments, along with BamHI/SalI-digested pNPTS139, were all ligated together using the NEB Hi-Fi DNA assembly kit according to the manufacturer’s instructions (New England Biolabs, Ipswich, MA). These plasmids were introduced into *B. melitensis* 16M, and merodiploid transformants were obtained by selection on Brucella agar plus 25 µg/ml kanamycin. A single kanamycin-resistant clone was grown overnight in Brucella broth and then plated onto brucella agar supplemented with 10% sucrose. Genomic DNA from sucrose-resistant, kanamycin-sensitive colonies was isolated and screened by PCR for replacement of the gene of interest.

### Mice

All animal studies were conducted in compliance with the University of Missouri Animal Care and Use Committee guidelines. We utilized 6- to 12-week-old C57BL/6J mice that were age and sex-matched for experiments. Mice were infected with 1×10^5^ CFUs of *B. melitensis* in 200 µL sterile PBS (sPBS) intraperitoneally (i.p.). For coinfections, a 1:1 ratio of WT *B. melitensis* 16M and a chloramphenicol resistant strain (*B. melitensisΔbmeI0265*) was prepared. Following infection, animals were maintained in individually ventilated caging under high-efficiency particulate air-filtered barrier conditions with 12-hour light and dark cycles within animal biosafety level 3 (ABSL-3) facilities at the University of Missouri. Mice were provided food and water ad libitum.

### Calculation of tissue bacterial burdens

Animals were euthanized, and spleens, livers, and reproductive tracts were harvested. Tissues were homogenized mechanically in sPBS (24). Serial dilutions were performed in triplicate in sPBS and plated onto brucella agar. Plates were incubated for three days at 37°C/5% CO_2_, colonies were counted, and the number of CFU/tissue or CFU/mL were calculated. For co-infection experiments, bacterial burdens were determined by plating homogenized spleens or cells on Brucella agar with or without 5 µg/mL chloramphenicol to select for *B. melitensisΔbmeI0265*.

### Flow cytometry

Spleens were homogenized and cell suspensions filtered through sterile 40 μm mesh following red blood cell lysis. Fc receptors were blocked in fluorescence-activated cell-sorting (FACS) buffer (2% heat inactivated fetal bovine serum in PBS) before extracellular staining with fluorochrome-conjugated mAbs from eBioscience or Biolegend (San Diego, CA) F4/80 (BM8), Ly-6G (1A8), CD11b (M1/70), and Ly-6C (HK1.4). Cells were then fixed in 4% paraformaldehyde at 4°C overnight before washing and resuspension in FACS buffer. Fluorescence was acquired on a CyAn ADP analyzer (Beckman Coulter, Brea, CA) and FlowJo (Tree Star, Ashland, OR) software was used for analysis. Cells were gated as Macrophages (F4/80^+^), monocytes (CD11b^+^Ly6C^high^), and neutrophils (CD11b^+^Ly6C^mid^).

### Metabolite quantification by gas chromatography-tandem mass spectrometry (GC-MS)

Nontargeted metabolomics was performed at the University of Missouri Metabolomics Center (25). ∼40 mg of spleen, liver, and reproductive tract was collected in a final concentration of 80% methanol. Tissues were homogenized and transferred to glass vials and incubated at room temperature with shaking (∼140 rpm) for 2 hours. Next, 1.5 mL of CHCl_3_ containing 10 µg/mL of docosanol (nonpolar internal standard) was added. The material was then sonicated, vortexed, and incubated at 50°C for 1 h. Then, 1 mL of HPLC grade H_2_O containing 25 µg/mL of ribitol (polar phase internal standard) was added, vortexed, and samples were incubated for 1 h at 50°C. Samples were centrifuged at 10°C at 3000x g for 40 minutes to pellet cell debris and separate phases. Upper phase (polar) and lower phase (nonpolar) were individually transferred to new glass tubes and dried in a speed vacuum. Material was stored at -20°C until ready to derivatize. For polar derivatization, samples were resuspended in a solution containing 50 µl of pyridine containing fresh 15 mg/ml methoxyamine-HCl, sonicated, vortexed, and placed in a 50°C oven for 1 h. Samples were allowed to equilibrate to room temperature and then 50 μl of N-methyltrimethylsilyltrifluoroacetamide (MSTFA) + 1% trimethylchlorosilane (TMCS) (Fisher Scientific) was added. Samples were vortexed, incubated for 1 h at 50°C, centrifuged, and transferred to glass inserts for injection. For nonpolar derivatization, samples were resuspended in 0.8 mL of CHCl_3_, 0.5mL of 1.25 M HCl in MeOH, vortexed and incubated for 4 h at 50°C. Post incubation, 2 mL of hexane was added to the samples, which were vortexed, and the upper layer was transferred to a new autosampler vial to dry. Material was then resuspended by adding 70 µl of pyridine, vortexed, and then 30 μl of MSTFA + 1%TMCS was added prior to incubation for 1 h at 50°C, centrifugation, and transfer to glass inserts for injection. Samples were analyzed using GC-MS on an Agilent 6890 GC coupled to a 5973N MSD mass spectrometer with a scan range from m/z 50 to 650. Separations were performed using a 60 m DB-5MS column (0.25-mm inner diameter, 0.25-mm film thickness; J&W Scientific) and a constant flow of 1.0ml/min helium gas. Results were interpreted using MetaboAnalyst 5.0 software (https://www.metaboanalyst.ca/) (26) with a P value threshold of 0.05.

### Macrophage generation, treatments, and infections

Cells were flushed from the femurs and tibias of C57BL/6J mice with sPBS supplemented with 5 µg/mL of gentamicin. Bone marrow-derived macrophages (BMDMs) were generated by cultured in complete medium with glutamine (CM; RPMI 1640, 10% fetal bovine serum [FBS], 10mM HEPES buffer, 10 mM nonessential amino acids, 10 mM sodium pyruvate) containing 30 ng/ml recombinant murine macrophage colony-stimulating factor (M-CSF; Shenandoah Biotechnology, Warwick, PA). After 3 days of culture, cells were washed with 30 mL of pre-warmed sPBS and fresh CM containing 30 ng/ml M-CSF was added to the culture flasks. After 3 days, adherent cells were collected by adding 0.05% trypsin (MilliporeSigma). Cells were plated at 1×10^6^ cells/ml in fresh CM (with or without glutamine) and allowed to adhere. Cells were infected at a multiplicity of infection (MOI) of 100 *B. melitensis* 16M or coinfected with 1:1 ratio of *B. melitensis* 16M and *B. melitensisΔbmeI0265* each at a MOI of 100. Cells were infected for 4 h, washed with sPBS, and then cultured in CM containing 50 μg/ml gentamicin for 30 minutes. Cells were then washed with sPBS and left to incubate in CM containing 2.5 μg/ml gentamicin for the remainder of the experiment. For GLS inhibition, 10 µm of Telaglenastat (MedChemExpress LLC, Monmouth Junction, NJ) was added to cells 12 h prior to infection (27–29). For GABAergic modulation, BMDMs were treated with GABA (100 µM) or bicuculline (BIC; 100 µM; GABA receptor antagonist) (Sigma-Aldrich, St. Louis, MO) at the same time that gentamicin containing media was added to the cells. At 24 h, 48 h, and 72 h post infection, supernatants were collected, and macrophages were washed and then lysed. These lysates were plated on brucella agar or brucella agar supplemented with chloramphenicol as explained above to determine the amount of intracellular *Brucella*. Supernatants were used for quantification of cytokines as described below.

### Cytokine quantification

Cell culture supernatants were filtered prior to measurement of cytokines. IL-1β levels were measured with a mouse IL-1β ELISA Ready Set Go kit (Invitrogen, Carlsbad, CA) according to the manufacturers’ instructions.

### Statistical analysis

For CFU and cytokines, data are expressed as mean +/- standard deviation (SD). Student’s unpaired T-test were used to assess differences in means between two groups with significance at p<0.05, while ANOVA followed by Tukey’s test with significance at p<0.05 was used for comparisons between ≥3 groups. N values and the number of experimental repeats are provided in the figure legends. All statistical analyses were performed with Prism software (version 10.1.1, GraphPad).

Multivariant statistical analysis for metabolite data was performed with Metaboanalyst (v5.0) (https://www.metaboanalyst.ca). The normalized values were used for statistical analyses such as principal component analysis (PCA), partial least squares discriminant analysis (PLS-DA), heatmaps, and volcano plots after log transformation and autoscaling with Metaboanalyst software. Pathway analysis was also performed in Metaboanalyst (V5.0) using KEGG database to identify the biological significance of metabolic pathways associated with infection. Pathways with P<0.05 were plotted indicating potential biological relevance.

## RESULTS

### Brucellosis progression in mice and tissue metabolism variations during different phases of infection

We investigated changes in tissue metabolite levels during progression of intraperitoneal *Brucella* infection in the spleens, livers, and female reproductive tracts of C57BL/6J mice. Bacterial loads peaked at day seven post-infection in all three tissues (Fig. 1A-C). At 14- and 28-days post-infection (dpi) CFU levels in spleens had a ∼5-10-fold reduction compared to 7 dpi (Fig. 1A). Similarly, in livers, there was a ∼100-fold reduction in CFUs between the 14 and 28 day timepoints relative to day 7 post-infection (Fig. 1B). In reproductive tracts, CFU counts were ∼10-fold lower at 14 dpi and ∼100-fold lower at 28 dpi compared to 7 dpi (Fig. 1C).

**Figure 1.**
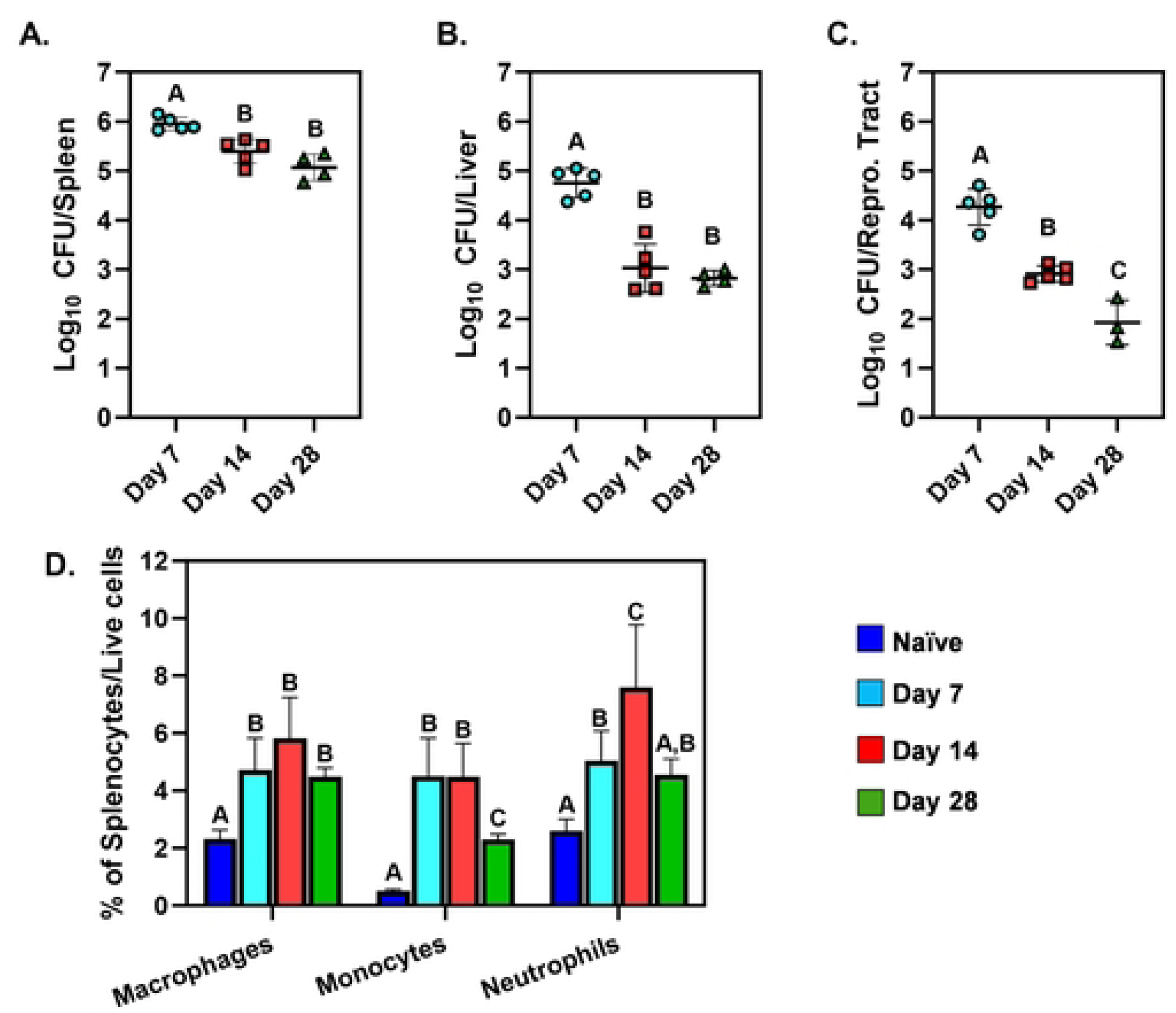
Brucellosis progression in mice. Spleens, livers, and reproductive tracts were harvested from naive C57BL/6J mice or from C57BL/6J mice at 7-, 14-, and 28-days after i.p. infection with 10^5^ CFUs of *B. melitensis* (n=4-6/group). Bacterial levels were determined in spleens **(A)**, livers **(B)**, and reproductive tracts **(C)**. Flow cytometry **(D)** was also performed to determine the percentages of macrophages (F4/80^+^), monocytes (CD11b^+^Ly6C^high^) and neutrophils (CD11b^+^Ly6C^mid^) amongst splenocytes. Error bars depict S.D. of the mean. Means with the same letter are not statistically different from each other (P<0.05 by ANOVA). Data are from one experiment.

Next, we analyzed the levels of innate immune cells in the spleen during infection. At 7 and 14 dpi the proportions of macrophages, monocytes, and neutrophils were significantly higher compared to naïve mice (Fig. 1D). At 14 dpi, neutrophil proportions were significantly increased compared to all other time points indicating that two weeks post-infection may be the peak of splenic inflammation in this model.

Based on these findings, we chose four time points to investigate metabolic changes during experimental brucellosis, pre-infection (naïve), 7 dpi representing the peak of bacterial loads, 14 dpi as the peak of inflammation, and 28 dpi as a starting point of chronic infection with some resolution of inflammation.

To investigate metabolic diversity within spleens, livers, and female reproductive tracts we extracted polar metabolites, which were then derivatized and analyzed via GC-MS. All three tissues demonstrated altered metabolic profiles when comparing infected animals with naïve mice. The clusters were plotted by Partial Least-Squares Discriminant Analysis (PLS-DA) (Fig. 2A-C) and the total numbers of compounds showing significant changes in spleens, livers, and reproductive tracts were 117, 30, and 32, respectively (Supplementary Tables 2.1-2.3)

**Figure 2.**
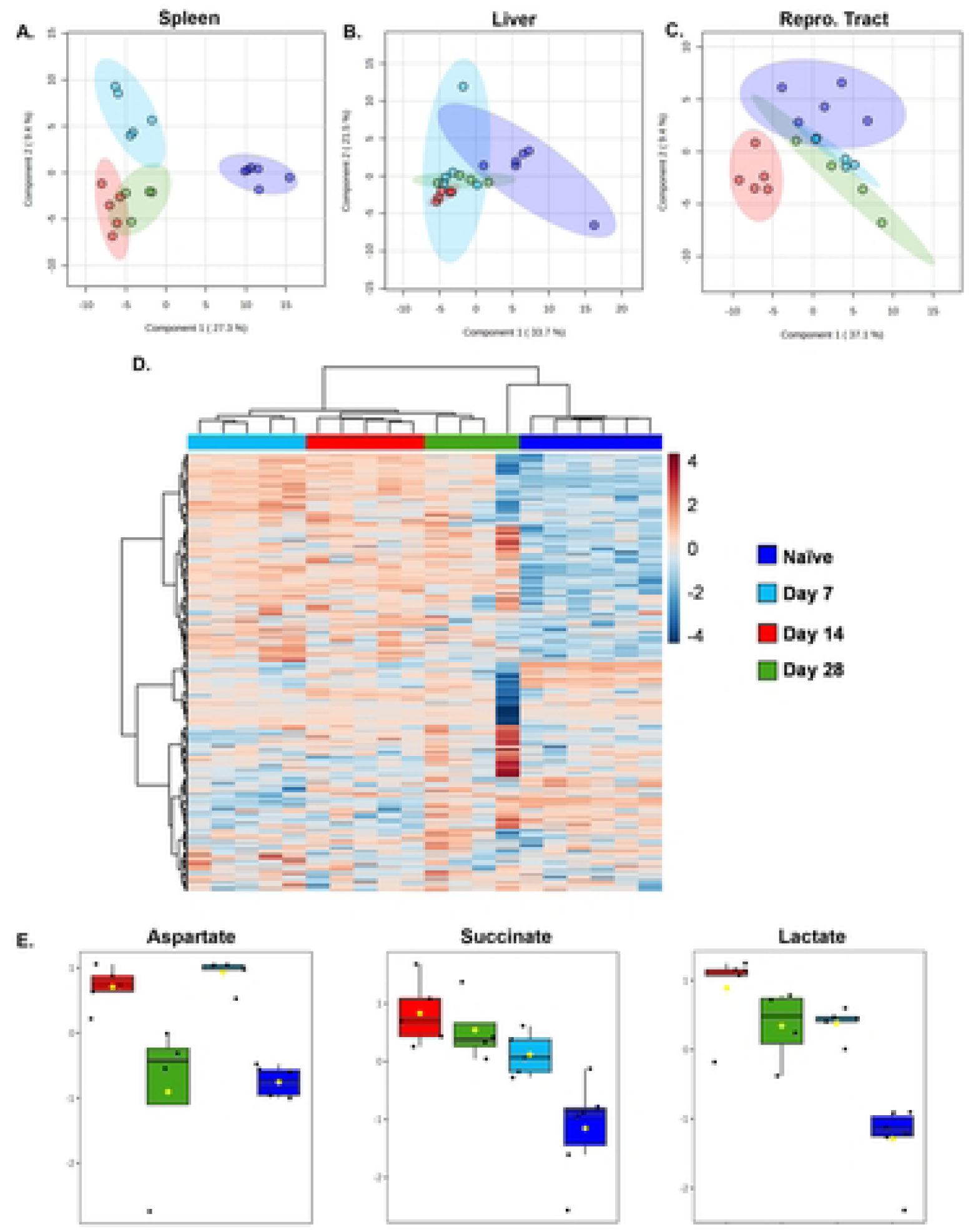
Tissue metabolism variations during different phases of *Brucella melitensis* infection using GCMS/MS analysis. C57BL/6J mice (WT, n=4-6 mice/group) were infected i.p. with 10^5^ CFUs of *B. melitensis*. Naive mice were included as controls. Spleens, livers, and reproductive tracts were harvested at 7-, 14-, and 28-days post-infection (dpi). Tissues were derivatized and analyzed via GC-MS. Score plot of Partial Least Square-Discriminant Analysis (PLS-DA) relative to time post-infection in spleens **(A)**, livers **(B)**, and female reproductive tracts **(C)**. Metabolite heatmap representation in spleen from tissues of naïve animals and from mice infected with *Brucella* for 7,14 or 28 days **(D)**. Box plots for select metabolites elevated at 14 dpi vs. naive mice determined by one-way ANOVA with Fisher’s LSD post hoc **(E)**. The Y axis is Log_10_ values of normalized instrument response for the select metabolites (aspartate, succinate, and lactate). Data are from one experiment.

Heat map visualization showed a distinct splenic metabolite profile separating uninfected from infected animals (Fig. 2D). The most remarkable changes in relative levels of compounds appeared to be between naïve and 14 dpi mice. We found elevated levels of aspartate, succinate, and lactate at 14 dpi in spleens (Fig. 2E). Lactate is the end product of glycolysis and therefore used as a marker for this pathway (18,30). Aspartate is associated with the urea cycle (31), and succinate was recently shown to be linked with HIF-1α stabilization and activation, which facilitates the metabolic shift from mitochondrial phosphorylation (OXPHOS) to glycolysis (19,32,33). Collectively, these findings from metabolite screening suggest a *Brucella-*driven change in tissue metabolism.

### Changes in tissue metabolism at the peak of Brucella-induced inflammation

To determine the metabolic profile of spleens, livers, and reproductive tracts at the peak of inflammation, we performed another metabolomic experiment on tissues from additional naive and 14 dpi animals. In this experiment, metabolite profiles in spleens and livers were again distinct in naïve and infected animals, clustering separately in the Principal Component Analysis (PCA). In contrast, reproductive tracts did not show defined separation, perhaps due to the absence of estrus synchronization before the experiment (Fig. 3A-C and Supplementary Tables 3.1-3.3). When compared to uninfected tissues, 42 out of 329 detected polar metabolites in reproductive tracts were significantly altered (T-test P<0.05) by *Brucella* infection, while in the spleens and livers, 68/205, and 139/329 metabolites were changed, respectively.

**Figure 3.**
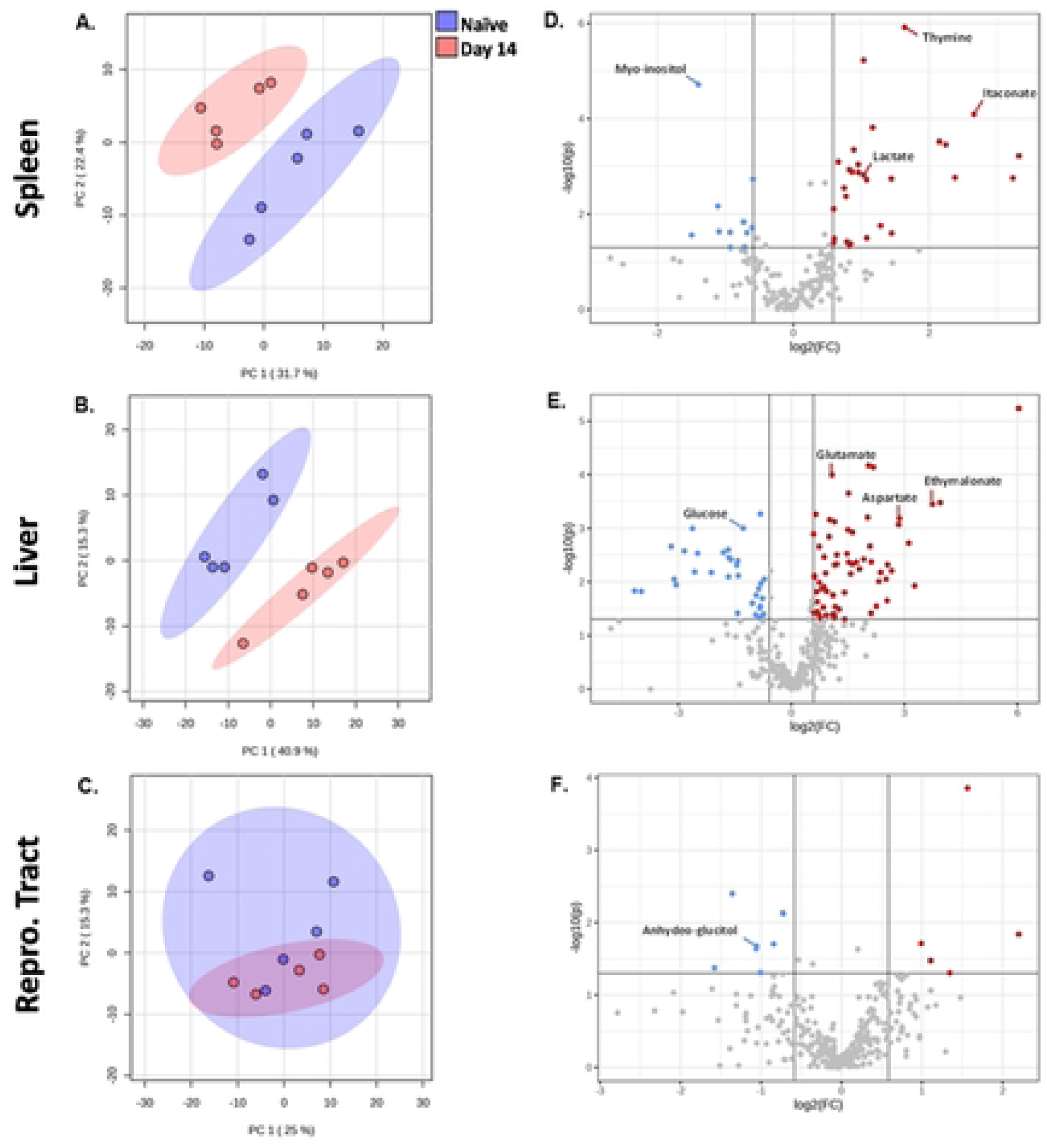
Global analysis of tissue metabolic profiles at the peak of inflammation after *Brucella melitensis* challenge. C57BL/6J mice (WT, n=5 mice/group) were infected i.p. with 10^5^ CFUs of *B. melitensis*. Principal Component Analysis (PCA) plot showing metabolic profiles at 14 dpi and in naïve mice, in spleens **(A)** livers **(B)**. and female reproductive tracts **(C)**. Volcano plot demonstrating down-(blue) and up-regulated (red) metabolites at 14 dpi compared with uninfected samples in spleens **(D)**, livers **(E)**, and female reproductive tract **(F)**. For each metabolite, the log2 fold change (threshold 1.5) is displayed on the x-axis and the -log10 (P-value) is displayed on the y-axis. P<0.05. Data are from one experiment.

We performed volcano tests combining fold change analysis (1.5 threshold) and T-test (P<0.05) to determine metabolites modified by infection (Fig. 3D-F). In spleens 11 compounds were lower in infected compared to naïve tissues, including myo-inositol (Fig. 3D). Twenty-nine compounds were significantly higher in infected spleens, including lactate, aspartate, itaconate, malate, and glutamate, suggesting an increase in the level of the TCA cycle intermediates in infected spleens (Fig. 4A). Livers from infected mice were found to have 34 compounds with lower relative levels and 67 compounds with significantly higher levels (Fig. 3B&E). Infected livers showed elevated pyroglutamate, aspartate, boric acid, and glutamate, among other metabolites (Fig. 4C). Five compounds were elevated, and 8 compounds were reduced in reproductive tracts from infected mice. Of these compounds, only 1,5-anhydroglucitol was identifiable (Fig. 3F).

**Figure 4.**
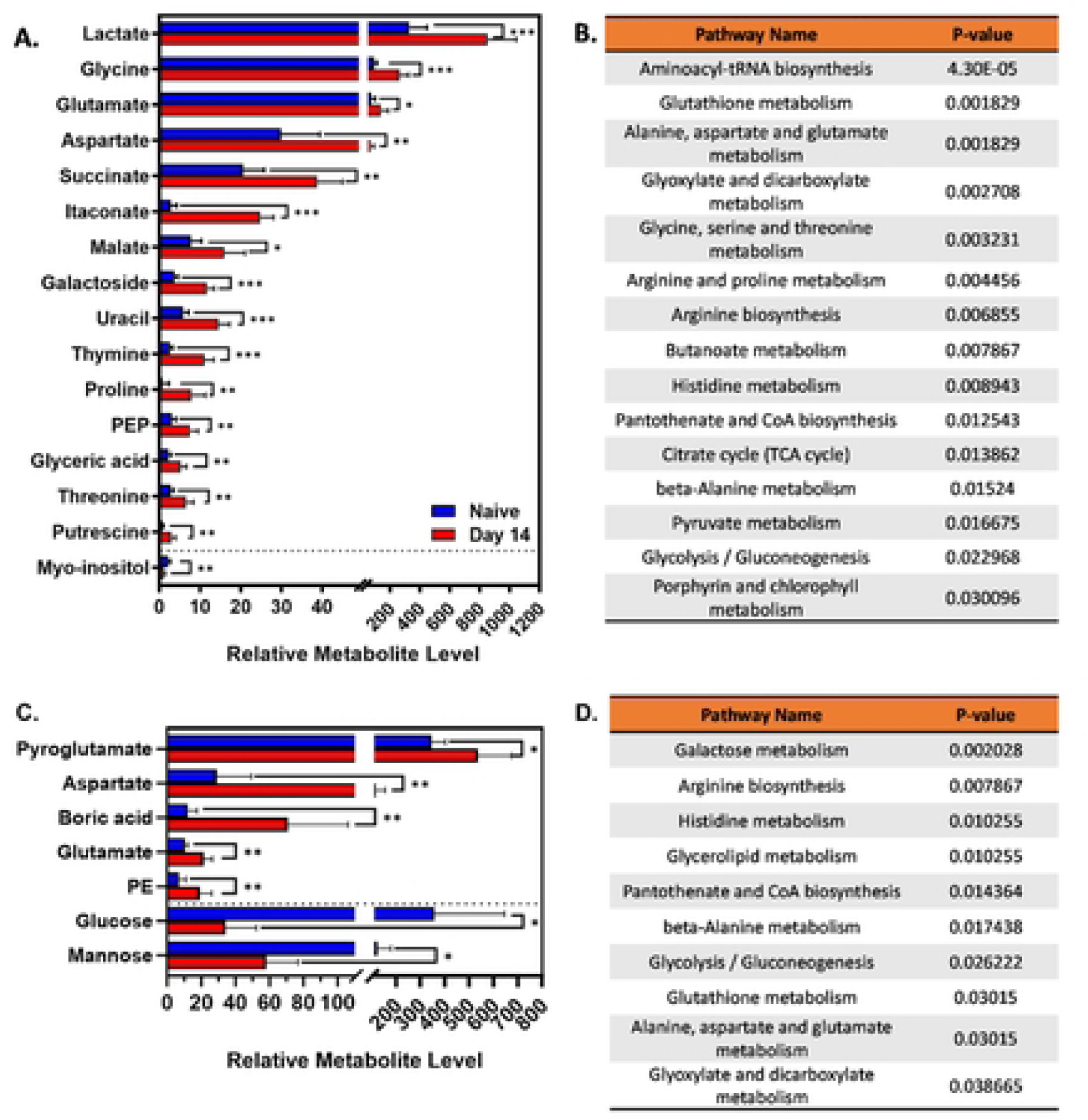
Peak of inflammation is associated with remarkable changes in tissue metabolic pathways. C57BL/6J mice (WT, n=5 mice/group) were infected i.p. with 10^5^ CFUs of *B. melitensis*. Relative metabolite levels and metabolic pathway analyses between 14 dpi and naïve mice in spleens **(A, B)** and livers **(C, D)** by GCMS/MS. Error bars stand for S.D. of the mean. * P<0.05; ** P<0.01; *** P<0.001 via T test. Data are from one experiment, n = 5 mice/group.

In addition, we performed a metabolic pathway analysis in spleens and livers targeting potential cellular signaling and metabolic networks that could play a role in *Brucella* infection. Several identified pathways modified by infection were linked to the TCA cycle, including glutaminolysis, the arginosuccinate shunt, glycolysis, and the GABA shunt (Fig. 4B&D). These data demonstrate a change in host metabolism concurrent with the peak of inflammation during infection. Of the identified pathways, there were some that might be involved with the cellular immune response against *Brucella*, including glutaminolysis and glycolysis.

### Inhibition of the glutaminolysis pathway dampens IL-1β production in response to Brucella

Our metabolomics data indicated alteration of the glutaminolysis pathway in response to *Brucella* infection. Glutamine is a vital compound in cellular metabolism and is involved in many functions, ranging from protein biosynthesis to mitochondrial respiration (34). Glutaminolysis is one of the means responsible for replenishing the TCA cycle via the breakdown of glutamine into glutamate. The key enzyme in that process is glutaminase (GLS). These reactions replenish the TCA cycle leading to the production of itaconate and succinate, which subsequently promotes a similar effect as the Warburg effect (aerobic glycolysis) in cancer cells (33). Therefore, we investigated the role of glutamine catabolism in BMDMs infected with *B. melitensis*.

To chemically inhibit the glutaminolysis pathway, we treated BMDMs with the GLS inhibitor CB-839, commercially known as Telaglenastat. GLS inhibition did not significantly affect the ability of macrophages to control intracellular *Brucella* infection at 24, 48, and 72 hours (Fig. 5A). However, Telaglenastat treatment dampened IL1-β secretion at 48 and 72 hours after infection (Fig. 5B). Glutamine-dependent anaplerosis is the largest source for succinate as a metabolite which in turn enhances IL1-β production (32). As glutamine is required for glutaminolysis and subsequent anaplerosis of the TCA cycle, we infected BMDMs with *B. melitensis* in complete media with and without glutamine. Despite bacterial clearance not being affected by glutamine, IL-1β secretion was dampened in the absence of glutamine 72 hours after infection (Fig. 5D). Collectively, these data demonstrate that glutaminolysis plays a role in IL-1β production but does not contribute to control of *Brucella* infection in BMDMs.

**Figure 5.**
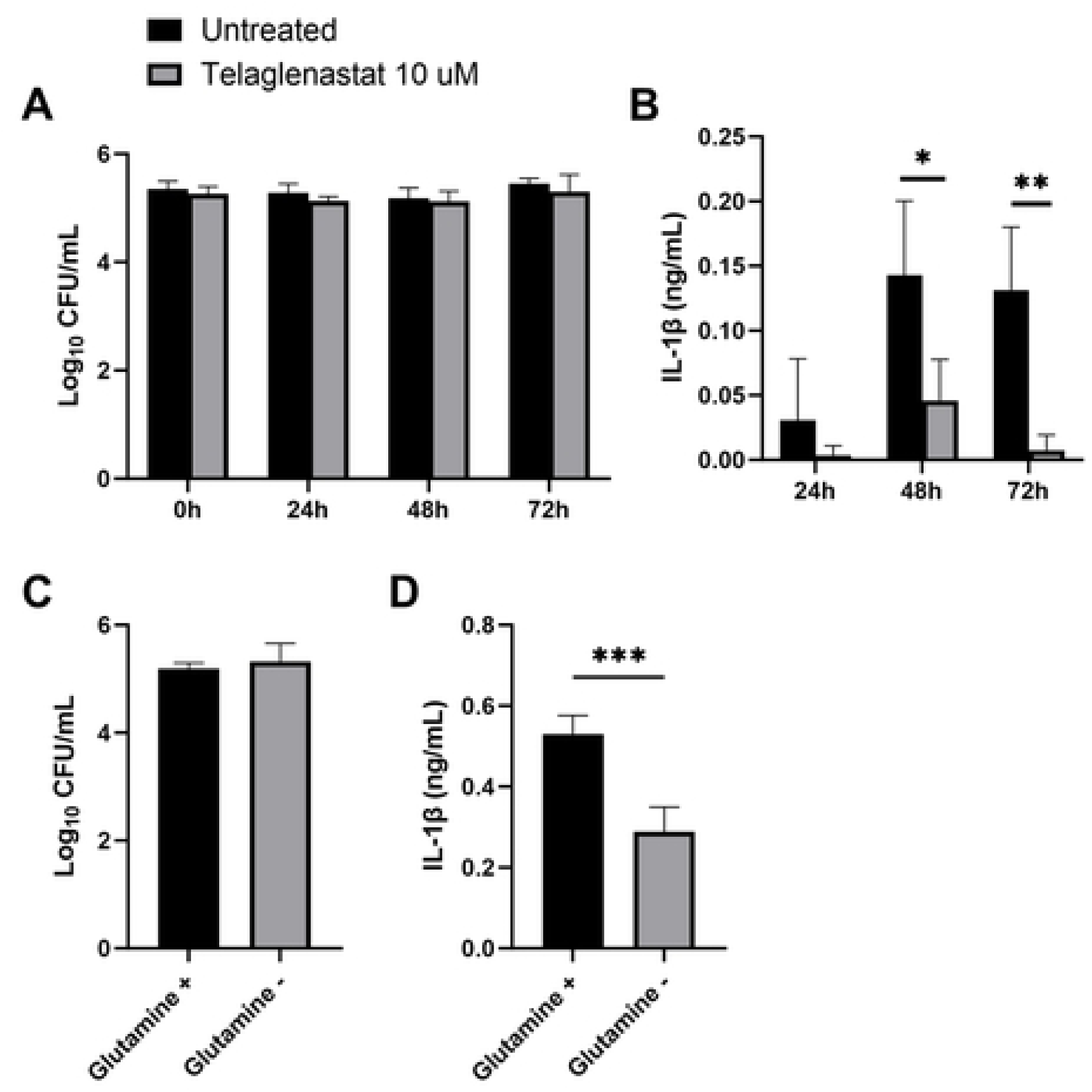
GLS inhibition decreases IL-1β secretion *in vitro*. Macrophages from C57BL/6J mice were infected with *B. melitensis* 16M at an MOI of 100, and at 0, 24, 48 and 72 hours post infection, intracellular CFU levels were determined **(A)** and IL-1β was measured in supernatants via ELISA **(B)**. Macrophages were infected with *B. melitensis* 16M at an MOI of 100 and cultured in media with or without glutamine supplementation, and intracellular bacteria were quantified **(C)** and IL-1β was measured in supernatants **(D)** 72 hours after infection. Error bars depict SD of the mean, n = 4 wells/group. Data are representative of 2 independent experiments. * P<0.05; ** P<0.01; *** P<0.001 compared to WT macrophages via T test.

### GABA supplementation does not alter control of *Brucella* by macrophages

Glutamate is also the precursor of GABA, an inhibitory neurotransmitter with the potential to enhance antimicrobial defenses against intracellular bacteria (35,36). To investigate if the host GABAergic system controls intracellular *B. melitensis* we treated infected BMDMs with GABA (100 µM) or bicuculline (BIC; 100 µM; GABA receptor antagonist) (Fig. 6A). No difference was observed between treatment groups at 48 hours.

**Figure 6.**
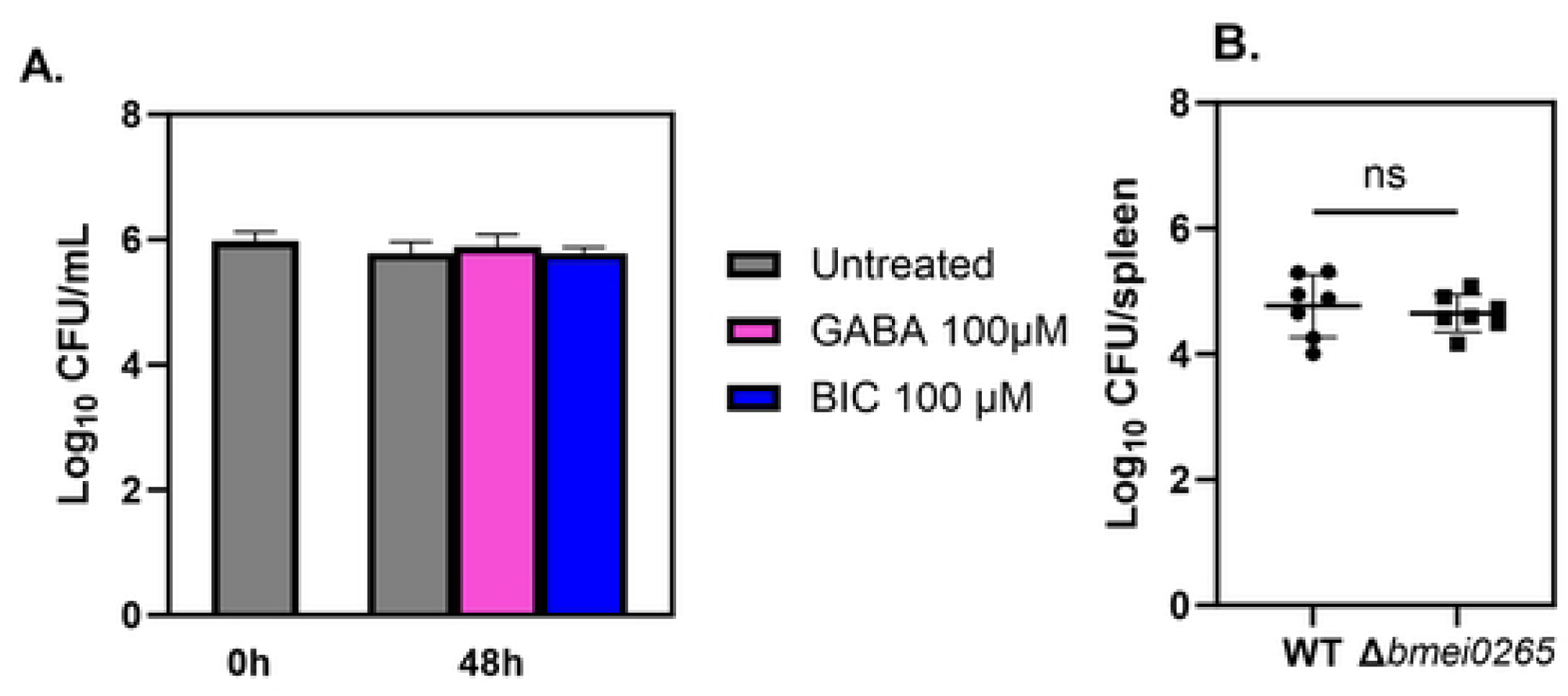
GABAergic system does not play a role in controlling *B. melitensis* 16M in BMDMs. **(A)** Macrophages from C57BL/6J mice (n =4 wells/group) were infected with *B. melitensis* 16M at an MOI of 100. After infection, cells were treated with GABA (100 µM) or bicuculline (BIC; 100 µM; GABA_A_R antagonist) and intracellular CFUs were determined at 0, 48 and 72 hours post-infection. **(B)** C57BL/6J mice (n=7) were challenged i.p. with a 1:1 ratio of 10^5^ *B. melitensis* 16M and *B. melitensisΔbmei0265*. and relative splenic CFU levels were determined fourteen days post-infection. Error bars depict SD of the mean. Data in **(A)** are representative of 2 independent experiments while data in **(B)** are combined from two experiments.

Our *in vivo* metabolomics data indicated that the GABA shunt was altered in mouse spleens at two-weeks post-infection (Fig. 4A&B). *B. abortus* has been previously shown to encode two GABA transporters, *bab1_1794* and *bab2_0879*, with moderate and high GABA import rates, respectively, but these GABA transporters were not required for virulence (37). There are other genes within the *Brucella* genome that are annotated to encode putative GABA transporters including *B. melitensis* BMEI0265. We therefore generated an isogenic BMEI0265 mutant (*B. melitensisΔbmei0265*). However, we did not find *B. melitensisΔbmei0265* to be attenuated in mouse spleens two weeks post-infection (Figure 6B). Although our findings suggest that *bmei0265* likely does not play a role in *B. melitensis* virulence, more studies are needed to understand the mechanisms of GABA signaling on host immune defense against *Brucella*.

## DISCUSSION

Over the past decade, immunometabolism studies have become key to understanding how metabolism of host cells impacts the outcome of infection (38). However, in the context of *Brucella* infection, the topic has not been extensively explored. Here, we investigated the interface of host-*Brucella* interactions by first screening tissue metabolic fluctuations during the course of infection. Untargeted GCMS/MS showed a *Brucella-*driven change in tissue metabolism at 14 dpi, particularly in metabolites correlated with the TCA cycle. Mitochondrial changes in metabolites related to the TCA cycle can regulate the function and activation of immune cells (39,40). We previously demonstrated the impact of itaconate, an intermediate metabolite from the TCA cycle, on mouse susceptibility to *Brucella* (18). In the present study, we found itaconate levels to be higher in spleens from infected mice (Fig. 4A). In line with these results, we suggest that TCA cycle-correlated metabolites have an impact on efficient immune responses against *B. melitensis*.

Metabolic pathway analysis displayed a link between infection and several pathways such as glutaminolysis, the arginosuccinate shunt, glycolysis, and the GABA shunt. Among the metabolites showing key changes, glutamate levels were significantly increased in infected livers compared to uninfected controls. Glutamate plays a role in macrophage polarization, replenishment of the TCA cycle via glutaminolysis, and in the GABA shunt at the mitochondria level (17,34,39,41). Glutaminolysis has been found to be critical in the metabolic reprogramming of M1-like human macrophages infected with *M. tuberculosis*, demonstrating its importance in the proinflammatory response (27). Similarly, we found that glutaminolysis inhibition with telaglenastat impairs secretion of the pro-inflammatory IL-1β cytokine (Fig. 5B). It is established that IL-1 is crucial to control *Brucella* infection in mice (42,43). However, while GLS inhibition dampened IL-1β production, we found that GLS inhibition did not affect the ability of BMDMs to control intracellular *B. melitensis* (Fig. 5A). Conversely, a study using a telaglenastat analog (BPTES) reported increased CFUs in a macrophage cell line infected with *B. abortus* (44). The divergent results could be due to differences between the two inhibitors, the types of macrophages, and the strains of *Brucella*.

Furthermore, glutamate is also involved in the GABA shunt, a process responsible for producing and conserving the supply of gamma-aminobutyric acid. GABA is an inhibitory neurotransmitter recently found to impact the host immune system (35,36,45). Studies performed with *M. tuberculosis*, *Salmonella* Typhimurium, and *Listeria monocytogenes* demonstrated that GABAergic system activation by GABA or its receptor agonist enhances macrophage antimicrobial defense against these intracellular bacteria (46). However, our results suggested that the GABAergic system does not play a role in controlling *B. melitensis* intracellularly.

The conversion of glutamate into GABA is catalyzed by the enzyme glutamate decarboxylase (GAD), which provides a pH homeostasis mechanism in some pathogenic bacteria, including *L. monocytogenes* (47). Several *Brucella* species have a functional GAD system; however, the system is lost in host-adapted pathogens such as *B. melitensis*, *B. abortus,* and *B. suis* (37). Nonetheless, *Brucella* has a GABA transporter with an undefined role in the metabolic utilization of GABA which may play a role in the pathogenesis of *Brucella* infection (37). According to our results, the potential transport of GABA transport by BMEI0265 does not play a role in *B. melitensis* virulence under the conditions tested here, however, further investigation would be needed to fully explore the role of GABA in the pathogenesis of brucellosis.

In conclusion, we show here that metabolite screening of spleens, livers, and reproductive tracts suggested a *Brucella*-driven change in tissue metabolism, with the most remarkable changes in host metabolism occurring at the peak of inflammation around two weeks after *B. melitensis* infection. Additionally, metabolite changes were related to intracellular pathways linked to mitochondria mechanisms. Between these candidate pathways, glutaminolysis was demonstrated to play a role in IL-1β production but did not contribute to macrophage control of *Brucella* infection *in vitro*.

## Acknowledgements

This work was supported by the National Institutes of Health (grants R01AI150797 and R21AI146397 to J. A. S.); and by funding from the University of Missouri Metabolomics Center.

## Supporting Information Captions

**Table S1:** Primers used in this study.

**Table S2.1:** Metabolite levels in spleens from naïve mice, and from spleens at 7, 14, and 28 days post-infection with *B. melitensis*.

**Table S2.2:** Metabolite levels in livers from naïve mice, and from livers at 7, 14, and 28 days post-infection with *B. melitensis*.

**Table S2.3:** Metabolite levels in reproductive tracts from naïve female mice, and from reproductive tracts at 7, 14, and 28 days post-infection with *B. melitensis*.

**Table S3.1:** Metabolite levels in spleens from naïve mice, and from spleens at 7 days post-infection with *B. melitensis*.

**Table S3.2:** Metabolite levels in livers from naïve mice, and from livers at 7 28 days post-infection with *B. melitensis*.

**Table S3.3:** Metabolite levels in reproductive tracts from naïve female mice, and from reproductive tracts at 7 days post-infection with *B. melitensis*.

## References

1. Gorvel JP, Moreno E. *Brucella* intracellular life: from invasion to intracellular replication. Vet Microbiol. 2002 Dec 20;90(1–4): 281–297.

2. Corbel MJ. Brucellosis: an overview. Emerg Infect Dis. 1997 Apr-Jun;3(2):213–21. doi: 10.3201/eid0302.970219.

3. Zhou C, Huang W, Xiang X, Qiu J, Xiao D, Yao N, et al. Outbreak of occupational Brucella infection caused by live attenuated *Brucella* vaccine in a biological products company in Chongqing, China, 2020. Emerg Microbes Infect. 2020;11(1):2544–2552. 10.1080/22221751.2022.2130099

4. Bossi P, Tegnell A, Baka A, Van Loock F, Hendriks J, Werner A, et al. Bichat guidelines for the clinical management of brucellosis and bioterrorism-related brucellosis. Euro Surveill. 2004;9(12):33–34. doi: 10.2807/esm.09.12.00506-en.

5. Zinsstag J, Roth F, Orkhon D, Chimed-Ochir G, Nansalmaa M, Kolar J, et al. A model of animal-human brucellosis transmission in Mongolia. Prev Vet Med. 2005 Jun;69(1–2):77–95. doi: 10.1016/j.prevetmed.2005.01.017.

6. Djokic V, Freddi L, de Massis F, Lahti E, van den Esker MH, Whatmore A, et al. The emergence of *Brucella canis* as a public health threat in Europe: what we know and what we need to learn. Emerg Microbes Infect. 2023 Dec;12(2): 224926. doi:0.080/222275.2023.2249126.

7. Yagupsky P, Morat P, Colmenero JD. Laboratory diagnosis of human brucellosis. Clin Microbiol Rev. 2019 Nov;33(1):e00073. doi: 10.1128/CMR.00073-19.

8. González-Espinoza G, Arce-Gorvel V, Mémet S, Gorvel J pierre. *Brucella*: reservoirs and niches in animals and humans. Pathogens. 2021 Feb 9;10(2):186. doi:10.3390/pathogens10020186.

9. Roop RM, Bellaire BH, Valderas MW, Cardelli JA. Adaptation of the brucellae to their intracellular niche. Mol Microbiol. 2004 May;52(3):621–30. doi: 10.1111/j.1365-2958.2004.04017.x.

10. Razei A, Javanbakht M, Hajizade A, Heiat M, Zhao S. Biomedicine & Pharmacotherapy Nano and microparticle drug delivery systems for the treatment of *Brucella* infections. Biomed Pharmacother. 2023 Dec;169:115875. doi:10.1016/j.biopha.2023.115875

11. Duran S, Anwar J, Moin ST. Interaction of gentamicin and gentamicin-AOT with poly-(lactide-co-glycolate) in a drug delivery system - density functional theory calculations and molecular dynamics simulation. Biophys Chem. 2023 Mar:294:106958. doi: 10.1016/j.bpc.2023.106958.

12. Gleeson LE, Sheedy FJ, Palsson-McDermott EM, Triglia D, O’Leary SM, O’Sullivan MP, et al. Cutting Edge: *Mycobacterium tuberculosis* Induces aerobic glycolysis in human alveolar macrophages that Is required for control of intracellular bacillary replication. J Immunol. 2016 Mar 15;196(6):2444–9. doi: 10.4049/jimmunol.1501612.

13. Zhang W, Wang G, Xu ZG, Tu H, Hu F, Dai J, et al. Lactate is a natural suppressor of RLR signaling by targeting MAVS. Cell. 2019 Jun 27;178(1):176–189.e15. doi: 10.1016/j.cell.2019.05.003.

14. O’Neill LAJ, Artyomov MN. Itaconate: the poster child of metabolic reprogramming in macrophage function. Nat Rev Immunol. 2019 May;19(5):273–281. doi: 10.1038/s41577-019-0128-5.

15. Barbier T, Zúñiga-Ripa A, Moussa S, Plovier H, Sternon JF, Lázaro-Antón L, et al. *Brucella* central carbon metabolism: an update. Crit Rev Microbiol. 2018;44(2):182–211. doi: 10.1080/1040841X.2017.1332002.

16. Liu Y, Xu R, Gu H, Zhang E, Qu J, Cao W, et al. Metabolic reprogramming in macrophage responses. Biomark Res. 2021 Jan 6: 9(1):1. doi: 10.1186/s40364-020-00251-y

17. Jha AK, Huang SCC, Sergushichev A, Lampropoulou V, Ivanova Y, Loginicheva E, et al. Network integration of parallel metabolic and transcriptional data reveals metabolic modules that regulate macrophage polarization. Immunity. 2015 Mar 17;42(3):419–30. doi: 10.1016/j.immuni.2015.02.005.

18. Lacey CA, Ponzilacqua-Silva B, Chambers CA, Dadelahi AS, Skyberg JA. MyD88-dependent glucose restriction and itaconate production control *Brucella* Infection. Infect Immun. 2021 Sep 16;89(10):e0015621. doi: 10.1128/IAI.00156-21.

19. Gomes MTR, Guimarães ES, Marinho F V., Macedo I, Aguiar ERGR, Barber GN, et al. STING regulates metabolic reprogramming in macrophages via HIF-1α during *Brucella* infection. PLoS Pathog. 2021 May 14;17(5):e1009597. doi: 10.1371/journal.ppat.1009597.

20. Xavier MN, Winter MG, Spees AM, Den Hartigh AB, Nguyen K, Roux CM, et al. PPARγ-mediated increase in glucose availability sustains chronic *Brucella abortus* infection in alternatively activated macrophages. Cell Host Microbe. 2013 Aug 14;14(2):159–70. doi: 10.1016/j.chom.2013.07.009.

21. Russell DG, Huang L, VanderVen BC. Immunometabolism at the interface between macrophages and pathogens. Nat Rev Immunol. 2019 May;19(5) :291–304. doi: 10.1038/s41577-019-0124-9.

22. Datsenko KA, Wanner BL. One-step inactivation of chromosomal genes in *Escherichia coli* K-12 using PCR products. Proc Natl Acad Sci U S A. 2000 Jun 6;97(12):6640–5. doi: 10.1073/pnas.120163297.

23. England JC, Perchuk BS, Laub MT, Gober JW. Global regulation of gene expression and cell differentiation in *Caulobacter crescentus* in response to nutrient availability. J Bacteriol. 2009 Nov 30;192(3):819–33. doi:10.1128/jb.01240-09.

24. Lacey CA, Mitchell WJ, Brown CR, Skyberg JA. Temporal role for MyD88 in a model of *Brucella*-induced arthritis and musculoskeletal inflammation. Infect Immun. 2017 Feb 23;85(3) :e00961–16. doi: 10.1128/IAI.00961-16.

25. Ponzilacqua-Silva B, Dadelahi AS, Abushahba MFN, Moley CR, Skyberg JA. Vaccine-elicited antibodies restrict glucose availability to control *Brucella* infection. J Infect Dis. 2024 Oct 16;230(4):e818–23. doi:10.1093/infdis/jiae172

26. Xia J, Psychogios N, Young N, Wishart DS. Metaboanalyst: A web server for metabolomic data analysis and interpretation. Nucleic Acids Res. 2009;37:W652–660.

27. Jiang Q, Qiu Y, Kurland IJ, Drlica K, Subbian S, Tyagi S, et al. Glutamine Is required for M1-like polarization of macrophages in response to *Mycobacterium tuberculosis* infection. mBio. 2022 Aug 30;13(4):e0127422. doi: 10.1128/mbio.01274-22.

28. Lampa M, Arlt H, He T, Ospina B, Reeves J, Zhang B, et al. Glutaminase is essential for the growth of triple-negative breast cancer cells with a deregulated glutamine metabolism pathway and its suppression synergizes with mTOR inhibition. PLoS One. 2017 Sep 26;12(9):e0185092. doi: 10.1371/journal.pone.0185092.

29. Bott AJ, Shen J, Tonelli C, Zhan L, Sivaram N, Jiang YP, et al. Glutamine anabolism plays a critical role in pancreatic cancer by coupling carbon and nitrogen metabolism. Cell Rep. 2019;29(5):1287–1298.e6. doi: 10.1016/j.celrep.2019.09.056

30. Rogatzki MJ, Ferguson BS, Goodwin ML, Gladden LB, Brooks GA. Lactate is always the end product of glycolysis. Front Neurosci. 2015 Feb 27:9:22. doi: 10.3389/fnins.2015.00022.

31. Allen AE, Dupont CL, Oborník M, Horák A, Nunes-Nesi A, Mccrow JP, et al. Evolution and metabolic significance of the urea cycle in photosynthetic diatoms. Nature. 2011 May 12;473(7346):203–7. doi: 10.1038/nature10074.

32. Tannahill GM, Curtis AM, Adamik J, Palsson-Mcdermott EM, McGettrick AF, Goel G, et al. Succinate is an inflammatory signal that induces IL-1β through HIF-1α. Nature. 2013 Apr 11;496(7444):238–42. doi: 10.1038/nature11986.

33. Li T, Copeland C, Le A. Glutamine Metabolism in Cancer. In: Advances in Experimental Medicine and Biology. Springer, Cham; 2021. p. 17–38. 10.1007/978-3-030-65768-0_2

34. Tapiero H, Mathé G, Couvreur P, Tew KD. II. Glutamine and glutamate. Biomed Pharmacother. 2002 Nov 1;56(9):446–57. doi: 10.1016/s0753-3322(02)00285-8.

35. Bhandage AK, Barragan A. GABAergic signaling by cells of the immune system: more the rule than the exception. Cellular and Molecular Life Sciences. 2021 Aug;78(15):5667–5679. doi: 10.1007/s00018-021-03881-z.

36. Xia Y, He F, Wu X, Tan B, Chen S, Liao Y, et al. GABA transporter sustains IL-1β production in macrophages. Sci Adv. 2021 Apr 7;7(15):eabe9274. doi: 10.1126/sciadv.abe9274.

37. Budnick JA, Sheehan LM, Benton AH, Pitzer JE, Kang L, Michalak P, et al. Characterizing the transport and utilization of the neurotransmitter GABA in the bacterial pathogen Brucella abortus. PLoS One. 2020 Aug 26;15(8):e0237371. doi: 10.1371/journal.pone.0237371.

38. O’Neill LAJ, Pearce EJ. Immunometabolism governs dendritic cell and macrophage function. J Exp Med. 2016 Jan 11;213(1):15–23. doi: 10.1084/jem.20151570.

39. Mehta MM, Weinberg SE, Chandel NS. Mitochondrial control of immunity: beyond ATP. Nat Rev Immunol. 2017 Oct;17(10):608–620. doi: 10.1038/nri.2017.66.

40. Diskin C, Ryan TAJ, O’Neill LAJ. Modification of Proteins by Metabolites in Immunity. Immunity. 2021 Jan 12;54(1):19–31. doi: 10.1016/j.immuni.2020.09.014.

41. Cruzat V, Rogero MM, Keane KN, Curi R, Newsholme P. Glutamine: Metabolism and immune function, supplementation and clinical translation. Nutrients. MDPI AG; 2018 6049 2018 Oct 23;10(11):1564. doi: 10.3390/nu10111564.

42. Skyberg JA, Thornburg T, Kochetkova I, Layton W, Callis G, Rollins MF, et al. IFN-γ-deficient mice develop IL-1-dependent cutaneous and musculoskeletal inflammation during experimental brucellosis. J Leukoc Biol. 2012 Aug;92(2):375–387. doi: 10.1189/jlb.1211626

43. Hielpos MS, Fernández AG, Falivene J, Alonso Paiva IM, Muñoz González F, Ferrero MC, Campos PC, Vieira AT, Oliveira SC, Baldi PC. IL-1R and inflammasomes mediate early pulmonary protective mechanisms in respiratory *Brucella abortus* infection. Front Cell Infect Microbiol. 2018 Nov 5;8:391. doi: 10.3389/fcimb.2018.00391.

44. Zhao T, Zhang Z, Li Y, Sun Z, Liu L, Deng X, Guo J, Zhu D, Cao S, Chai Y, Nikolaevna UV, Maratbek S, Wang Z, Zhang H. *Brucella abortus* modulates macrophage polarization and inflammatory response by targeting glutaminases through the NF-κB signaling pathway. Front Immunol. 2023 May 31;14:1180837. doi: 10.3389/fimmu.2023.1180837.

45. Bhat R, Axtell R, Mitra A, Miranda M, Lock C, Tsien RW, Steinman L. Inhibitory role for GABA in autoimmune inflammation. Proc Natl Acad Sci U S A. 2010 Feb 9;107(6):2580–5. doi: 10.1073/pnas.0915139107.

46. Kim JK, Kim YS, Lee HM, Jin HS, Neupane C, Kim S, Lee SH, Min JJ, Sasai M, Jeong JH, Choe SK, Kim JM, Yamamoto M, Choy HE, Park JB, Jo EK. GABAergic signaling linked to autophagy enhances host protection against intracellular bacterial infections. Nat Commun. 2018 Oct 10;9(1):4184. doi: 10.1038/s41467-018-06487-5.

47. Ryan S, Hill C, Gahan CG. Acid stress responses in *Listeria monocytogenes*. Adv Appl Microbiol. 2008;65:67–91. doi: 10.1016/S0065-2164(08)00603-5.

